# Dermal macrophages set pain sensitivity by modulating tissue NGF levels through SNX25–Nrf2 signaling

**DOI:** 10.1101/2021.01.26.428327

**Authors:** Tatsuhide Tanaka, Hiroaki Okuda, Yuki Terada, Takeaki Shinjo, Mitsuko Banja, Kazuya Nishimura, Ayami Isonishi, Hidemasa Furue, Shoko Takemura, Kouko Tatsumi, Akio Wanaka

## Abstract

Crosstalk between peripheral neurons and immune cells plays important roles in pain sensation. We identified *sorting nexin 25* (*Snx25*) as a pain-modulating gene in a transgenic mouse line with reduced pain behavior. *Snx25* conditional-KO (cKO) in monocyte/macrophage-lineage cells but not in the peripheral sensory neurons reduced pain responses in both normal and neuropathic conditions. Cross transplantation experiments of bone marrows between cKO and wild type (WT) mice revealed that cKO macrophages caused dull phenotype in WT mice and WT macrophages in turn increased pain behavior in cKO mice. SNX25 in dermal macrophages enhances NGF (one of the key factors in pain sensation) production by inhibiting ubiquitin-mediated degradation of Nrf2, a transcription factor that activates *Ngf* mRNA synthesis. We conclude that dermal macrophages set pain sensitivity by producing and secreting NGF into the dermis in addition to their host defense functions.

## Introduction

The skin is frequently stressed by mechanical trauma. Sensory stimuli impinging on skin are encoded by peripheral sensory neurons that can be classified into low-threshold mechanoreceptors (LTMRs), which detect innocuous tactile stimuli, and nociceptors, which exclusively respond to harmful stimuli (Abraira and Ginty, 2013) (Basbaum et al., 2009). Dorsal root ganglion (DRG) neurons are highly diverse in terms of cell size, gene expression and myelination level. While small-diameter neurons are the pain-sensing neurons, medium- to large-diameter neurons preferentially detect low-threshold mechanical stimulation (Liu and Ma, 2011). Tissue damage of skin leads to the release of inflammatory mediators by activated nociceptors or by nonneural cells that reside within or infiltrate into the injured area, including macrophages, mast cells, neutrophils, keratinocytes, and fibroblasts. These inflammatory mediators, such as serotonin, histamine, glutamate, ATP, and nerve growth factor (NGF), act directly on the nociceptors. Although peripheral sensitization after tissue injury is well described in the skin (Basbaum et al., 2009), the roles of inflammatory cells under normal conditions or in acute pain sensation are not fully understood. Recent studies have uncovered a close association of macrophages with peripheral neurons; tissue macrophages can be divided into two subsets, namely a nerve-associated and a blood vessel-associated population (Chakarov et al., 2019). A subset of skin macrophages is closely associated with peripheral nerves and promotes their regeneration when damaged (Kolter et al., 2019). In neuropathic conditions, macrophages can accelerate pain sensation by sensing tissue angiotensin 2 (Shepherd et al., 2018a) or complement 5a (Shutov et al., 2016) (Warwick et al., 2019). In the latter model, macrophages activate a “vicious cycle”: macrophages secrete NGF into tissues and the NGF in turn stimulates macrophages.

NGF is important for generation of pain and in hyperalgesia in diverse pain states (Barker et al., 2020). In addition to enhancing the activity of nociceptive ion channels to promote rapid depolarization and sensitization, NGF also mediates changes in gene expression and membrane localization, both of which contribute to increased sensory neuron excitability (Mamet et al., 2003) (Stratiievska et al., 2018). In humans, hereditary sensory and autonomic neuropathy type V (HSAN V) (OMIM 608654), characterized by a marked absence of pain sensibility, is caused by mutations in the *Ngf* gene (Einarsdottir et al., 2004) (Capsoni, 2014). Mouse models for HSAN V have been generated, in which the biological activity of NGF is blocked either by neutralizing antibodies (Ruberti et al., 2000) or by *Ngf* expression being abolished with homologous recombination (Chen et al., 1997). These mice show a significant reduction of sensory innervations, which leads to decreased pain perception (Capsoni, 2014). Collectively, these studies show that NGF is expressed in immune cells including macrophages and facilitates pain transmission by sensory neurons through a variety of mechanisms. Although the level of NGF should be maintained within an optimal range for sensing the normal environment and for evading noxious pain sensation, the mechanisms underlying its regulation remain to be determined.

We serendipitously discovered a pain-insensitive transgenic mouse line. Forward genetic analyses of the mouse led us to identify *sorting nexin 25* (*Snx25*) as a pain-modulating gene. SNX family members are involved in membrane trafficking, cell signaling, membrane remodeling, and organelle motility (Cullen and Korswagen, 2012). We demonstrate here that SNX25 in dermal macrophages activates NGF production by inhibiting ubiquitin-mediated degradation of Nrf2, one of the key transcription factors that activates *Ngf* mRNA transcription (Mimura et al., 2011). SNX25 in dermal macrophages modulates acute pain sensing under both normal and painful conditions via NGF/TrkA signaling. These findings indicate that macrophage-to-neuron signaling is important in pain processing even in naïve skin in addition to in neuropathic or inflammatory situations.

## Results

### *Snx25*+/− mice show a pain-insensitive phenotype

We serendipitously found that pain responses to mechanical stimuli were reduced in Tg (Mlc1-tTA) #Rhn mice (strain name, B6; CBB6(129)-Tg (Mlc1-tTA) 2Rhn) during handling and genotyping of the mice (Tanaka et al., 2010) (**Figure 1A**). The TG mouse was on a mixed genetic background of the 129S6, CBA, and C57BL/6J strains. To negate the possibility that the pain-insensitive phenotype was derived from the mixed genetic background, we back-crossed the TG mice with C57BL/6J mice for seven generations to obtain a genetic background indistinguishable from that of wild type (WT) C57BL/6J mice. Even with the same genetic background, the pain response to mechanical stimuli was also reduced in the TG mice (**Figure 1B**). Notably, the TG mice were insensitive to mechanical stimuli in normal conditions without any neuropathic or inflammatory paradigm (**Figures 1A and 1B**). We also noticed that pain responses to a chemical stimulus (5% formalin injection), such as lifting, shaking, and licking of the paw, were significantly reduced in the TG mice (**Figures 1C and 1D**). Immunohistochemistry revealed that the number of c-Fos-positive cells in the spinal dorsal horn after 5% formalin injection into hind paw skin was lower in TG than in WT mice (**Figures S1A** and **S1B**). Since the TG mouse harbors a BAC transgene (clone RP23-114I6, 198kb), we first speculated that an exogenous gene(s) in the BAC might modulate pain behavior. Using next-generation sequencing (NGS), we determined the genome insertion site of the BAC transgene (83 kb out of 198 kb) into 8qB1.1 of chromosome 8 in the TG mouse (**Figure S1C**). The expression levels of exogenous BAC-borne *Mlc1*and *Mov10l1*, however, were indistinguishable from those in WT mice (**Figures S1C–S1E**). We next hypothesized that the transgene might affect endogenous gene expression and thereby influence pain behavior. NGS analyses also revealed that the transgene (83kb) was inserted in the 8qB1.1 region, resulting in deletion of three genes (*Snx25*, *Slc25a4*, and *Cfap97*) (**Figure S1C**). Subsequent cDNA microarray analyses confirmed that these endogenous gene expressions were almost null (**Figure S1F**). One or a combination of these gene knockouts could be responsible for the pain behavior. We focused on *Snx25* (**Figures S1G** and **H**) and obtained commercially available *Snx25* knockout (KO) mice (Nanjing BioMedical Research Institute of Nanjing University; strain name, B6/N-*Snx25*^tm1a/Nju^, strain number, T001400, https://www.mousephenotype.org/data/genes/MGI:2142610) and checked pain behavior in the KO mice. The KO construct was a KO-first conditional allele targeting vector allowing expression monitoring (Skarnes et al., 2011). In the targeting construct, an *En2SA*-IRES-*lacZ* cassette was inserted upstream of exon 4 of *Snx25*, to create a null allele by splicing and premature termination of the transcript (**Figure 2A**). SNX25 is widely expressed in different tissues with a particular abundance in the lung (Hao et al., 2011). *Snx25* −/− mice are embryonic lethal, and therefore we first measured SNX25 expression in the lung of heterozygotes. SNX25 expression in the *Snx25* +/− mouse was approximately half of that in the WT mouse (**Figure 2B**). The pain responses to mechanical and chemical stimuli were reduced in *Snx25* +/− mice to levels comparable to those in *Mlc*1 TG mice (**Figures 2C–2E**). Although thermal nociception in the *Snx25* +/− mice was not affected at 2 months of age, older mice of 6-8 months-old displayed a higher latency to respond to a heat stimulus (**Figure S2A**). To further determine whether the *Snx25* +/− mice show a dull phenotype after nerve injury, we assessed mechanical hypersensitivity induced by spared nerve injury (SNI) (Decosterd and Woolf, 2000). The responses of the *Snx25* +/− mice were significantly attenuated as compared to the WT mice after SNI (**Figure 2F**). These results indicate that pain responses were reduced not only under normal conditions but also under painful conditions in the *Snx25* +/− mice. We checked cellular size distribution and the expression of small (CGRP) and large (NF200) neuron markers in the DRG of the *Snx25* +/− mice. The sensory neurons of the heterozygotes were indistinguishable from those of the WT mice, indicating that abnormal reactions to pain stimuli were not the result of the loss of particular neuronal populations (**Figures S2B–D**).

**Figure 1.**
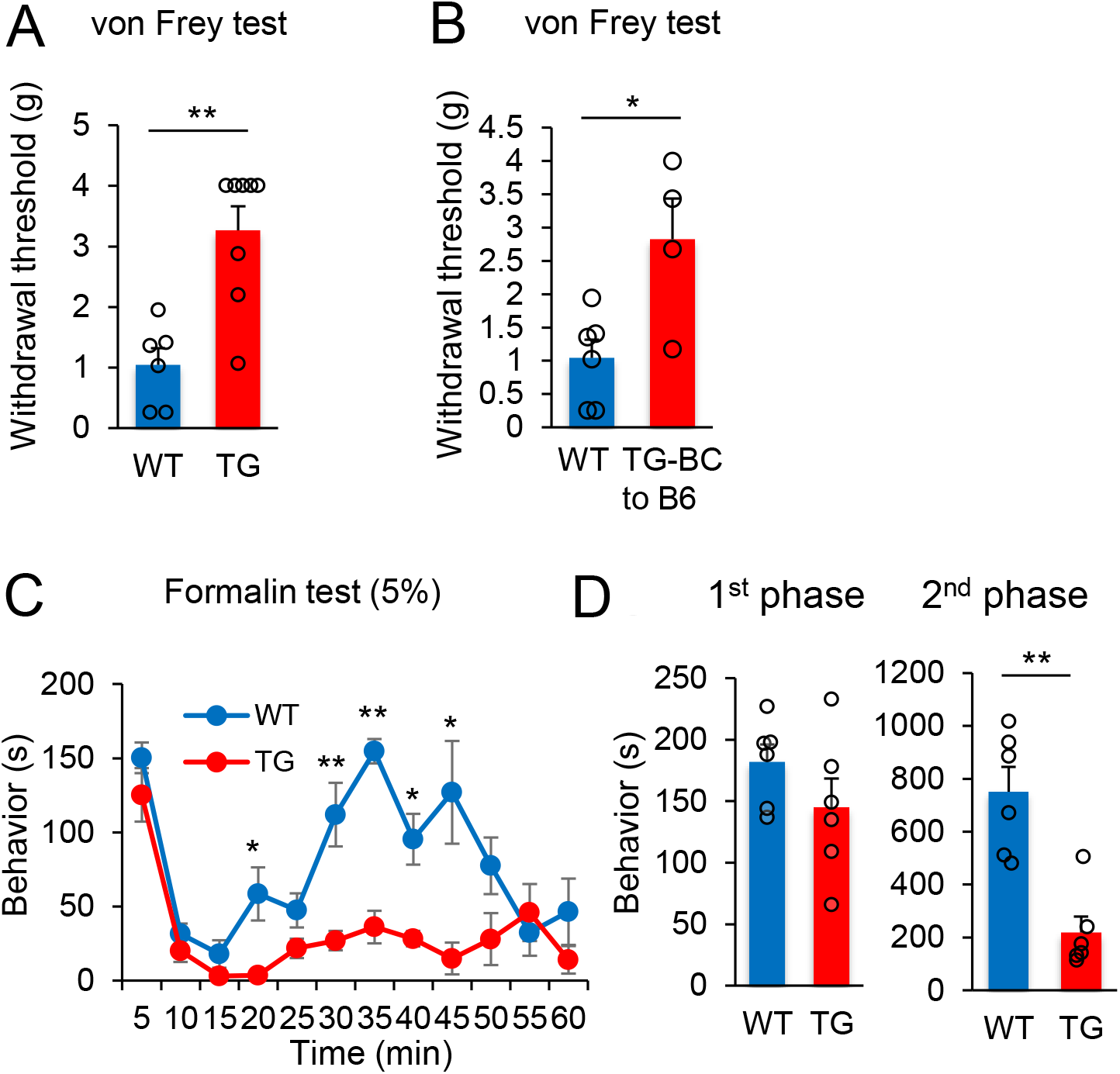
*Mlc1* TG mice show a pain-insensitive phenotype. (**A**) Comparison of paw withdrawal thresholds to mechanical stimulation with von Frey filaments between wild type (WT; n = 6) and *Mlc1* TG mice (TG; n = 8). (**B**) The same von Frey test except that *Mlc1* TG mice (mixed 129S6/CBA/C57BL/6J background) were backcrossed with C57BL/6J mice for 7 generations (WT: n = 6; Mlc-1 TG-BC to B6: n = 4). (**C**) Formalin test of wild type and *Mlc1* TG mice. Pain-related behavior time including licking, lifting, and flinching of the 5% formalin-injected paw was plotted for 5-minute periods (WT: n = 6; TG: n = 6). **(D)** Left, measurement of the behavior time in 1^st^ phase (0–10 min). Right, measurement of the behavior time in 2^nd^ phase (20–60 min). Results are represented as mean ± SEM of 3–5 independent experiments. Statistical significance was calculated using the Student’s *t*-test. \**p* < 0.05, \*\**p* < 0.01. See also Figure S1.

**Figure 2.**
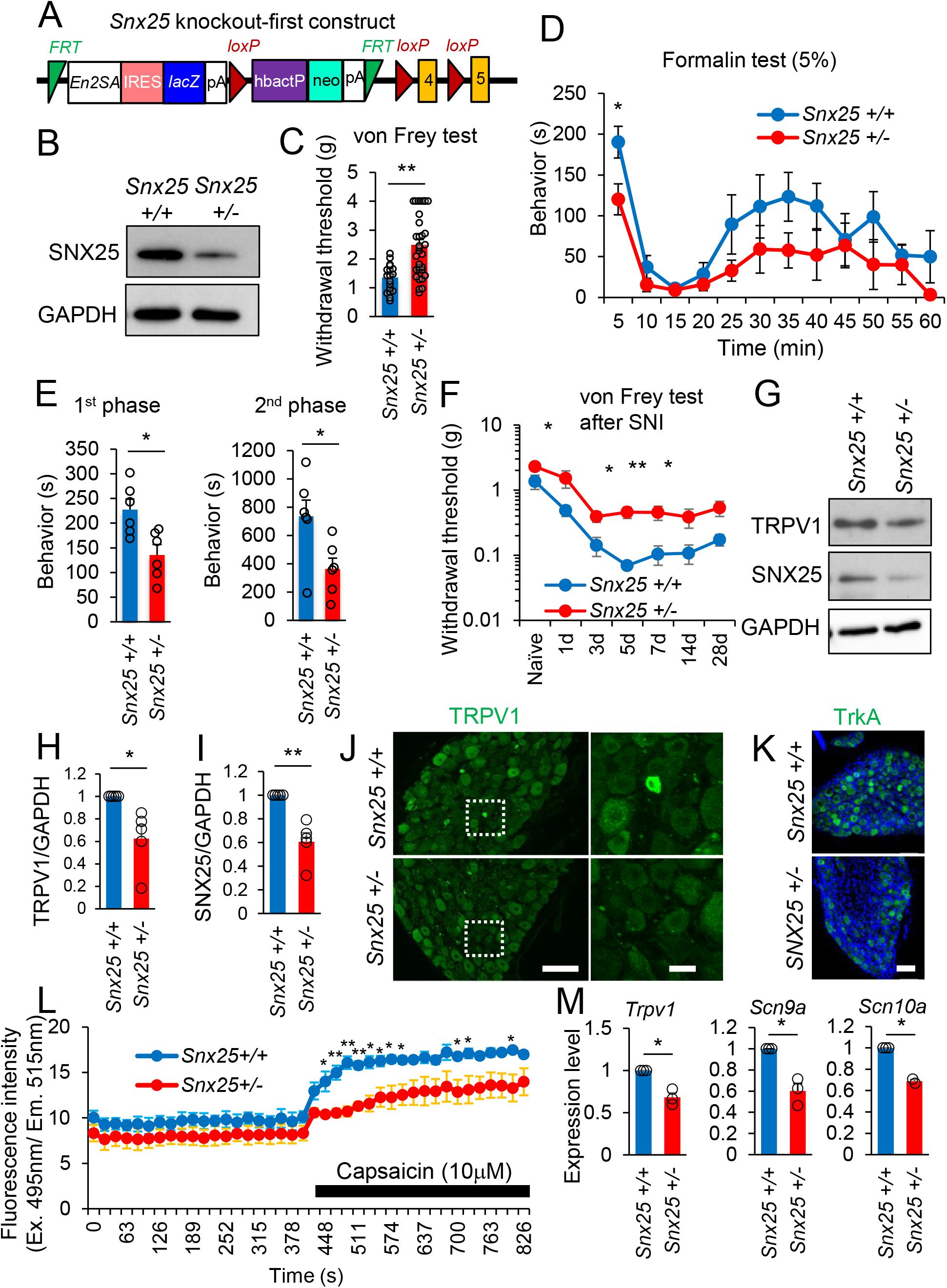
*Snx25* +/− mice show a pain-insensitive phenotype. (**A**) Scheme of the targeting vector used to knock out the *Snx25* gene. An *En2SA*-IRES-*LacZ* cassette was inserted upstream of exon 4. *FRT* and *loxP* sites enable the conditional deletion of gene segments. (**B**) Expression level of SNX25 in the lung of WT and *Snx25* +/− mice. (**C**) von Frey test showing significant elevation of withdrawal thresholds in heterozygotes relative to WT mice. (WT: n = 16; *Snx25* +/−: n = 28). (**D**) Formalin test of WT and *Snx25* +/− mice. Pain-related behavior time including licking, lifting, and flinching of the formalin (5%)-injected paw was plotted for 5-minute periods (WT: n = 6; *Snx25* +/−: n = 6). **(E)** Formalin test of wild type and heterozygote mice. Left panel shows total pain-related behavior time in 1^st^ phase (0–10 min). Right panel indicates 2^nd^ phase (20–60 min). Heterozygotes exhibited significantly shorter behavior times than WT mice in both 1^st^ and 2^nd^ phases. (**F**) Mechanical allodynia was evaluated by von Frey test after spared nerve injury (SNI) in mice (WT: n = 4; *Snx25* +/−: n = 7). *Snx25* heterozygote mice showed dull responses to von Frey filaments as compared to the WT mice. (**G**) Representative Western blots show expression levels of TRPV1 and SNX25 in the DRG (L4) of WT and *Snx25* +/− mice. (**H** and **I**) Semi-quantitative analyses of Western blotting data for TRPV1 (H) and SNX25 (I) (WT: n = 5; *Snx25* +/−: n = 5). (**J**) Confocal images of the DRG (L4) stained with anti-TRPV1 antibody in WT and *Snx25* +/− mice. Scale bar, 100 μm. Right panels show magnified views of boxed areas in the corresponding left panels. Scale bar, 20 μm. (**K**) Confocal images of the DRG (L4) of WT and *Snx25* +/− mice, stained with anti-TrkA antibody. Scale bar, 100 μm. (**L**) Fluo-4 Ca imaging of primary DRG neurons from an entire well (96-well plate) dissociated from WT and *Snx25* +/− mice (WT: n = 3; *Snx25* +/−: n = 3). Fluorescence intensities in response to 10 μM capsaicin treatment were significantly lower in the *Snx25* +/− DRG neurons. (**M**) mRNA expression levels for pain-related factors (*Trpv1*, *Scn9a*, *Scn10a*) in DRG (L4) of *Snx25* +/− mice were significantly lower than those of WT mice (WT: n = 3; *Snx25* +/−: n = 3). Results are represented as mean ± SEM of 3–5 independent experiments. Significance was calculated using the Student’s *t*-test (C, D, E and F) or Welch’s *t*-test (H, I and M). \**p* < 0.05, \*\**p* < 0.01. See also Figure S2.

### Pain-related factors are reduced in *Snx25* +/− mice

To further investigate the roles of SNX25 in pain sensation, we examined the expression of pain-related factors. Consistent with the pain-insensitive phenotype, the expression of pain-related factors including TRPV1 and TrkA was down-regulated in the DRG, sciatic nerve, and spinal cord of *Snx25* +/− mice (**Figures 2G–K, Figures S2E-F**). Capsaicin is a well characterized compound that produces a sensation of pain and stimulates TRPV1 channels on peripheral sensory nerves. Capsaicin elevated the intracellular Ca level in a population of primary cultured DRG neurons, but the amplitude of this Ca elevation was significantly lower in *Snx25* +/− neurons than in WT neurons, indicating that SNX25 deficiency resulted in TRPV1 channel inactivation in the DRG neurons (**Figure 2L**). We also observed that the mRNA levels of *Trpv1*, *Scn9a*, and *Scn10a*, which are related to pain perception, were reduced in *Snx25* +/− DRGs (Barker et al., 2020) (**Figure 2M**). From these data, we conclude that the pain-insensitive phenotype of the *Snx25*+/− mice was due to reduced levels of pain-related factors in the peripheral sensory neurons.

### DRG-specific *Snx25* cKO mice do not show pain-insensitive phenotype

To further define the tissues and cells responsible for the pain-insensitive phenotype in *Snx25*+/− mice, conditional alleles were generated by removal of the gene-trap cassette by Flippase (FLP), which reverts the mutation to wild type (WT), leaving *loxP* sites on either side of the critical exon 4 (Skarnes et al., 2011) (Yamazaki et al., 2016) (**Figure S3A**). We found that pain responses to mechanical and chemical stimuli reverted to the normal level in *Snx25^loxP/loxP^* mice (**Figures S3B and S3C**), underlining the idea that the pain-insensitive phenotype was due to the lack of the *Snx25* gene. To further assess the role of SNX25 in pain behavior, we next conditionally knocked out *Snx25* in the DRG by crossing *Snx25^loxP/loxP^* mice with *Advillin* (*Avil*)^*Cre*ERT2^ mice (Lau et al., 2011). We administered 0.05% tamoxifen (TAM) orally for 2 weeks (**Figure S4A**), a method that is convenient for continuous administration and results in efficient induction of recombination while minimizing stress on the mice (Kiermayer et al., 2007). Continuous feeding with TAM-containing chow markedly reduced expression of SNX25 in the DRG (**Figure S4B**). Contrary to our expectation, *Avil^Cre^*^ERT2/WT^; *Snx25^loxP/loxP^* mice had normal pain responses to both mechanical and chemical stimuli (**Figures S4C–S4F**). We also observed that the mRNA levels of *Trpv1*, *Scn9a*, and *Scn10a* were not reduced in DRGs of *Avil^Cre^*^ERT2/WT^; *Snx25^loxP/loxP^* mice (**Figure S4G**). These results indicate that SNX25 in the DRG neither regulates pain-related factors nor affects pain sensation.

### SNX25 in BM-derived macrophages contributes to pain sensation

Given that the DRG cKO showed normal pain behavior, we next focused on immune cells, since *Snx25* +/− mice showed the pain-insensitive phenotype in an acute inflammatory pain model (**Figures 2D and 2E**). Among immune cells in the skin, a subset of macrophages in the dermis are associated with peripheral nerves (Chakarov et al., 2019), and are therefore good candidates for pain-regulating cells. We confirmed that cells from a population of dermal macrophages (MHCII, CD206, and F4/80-positive; Chakarov et al.2019) were closely associated with PGP9.5-positive sensory fibers (**Figure 3A**) and were SNX25-immunoreactive (**Figure S5A**). We first checked whether SNX25 in these macrophages could influence migration from bone marrow to dermis. Immunohistochemistry revealed that the numbers of CD206- or MHCII-positive macrophages in hind paw skin in *Snx25* +/− mice were normal (**Figure S5B**). The expression level of CD206 in hind paw skin in *Snx25* +/− mice was also similar to that in *Snx25* +/− mice (**Figures S5C** and **D**). These results indicate that the pain-insensitive phenotype in the *Snx25*+/− mice is not due to reduced dermal macrophages under normal conditions.

**Figure 3.**
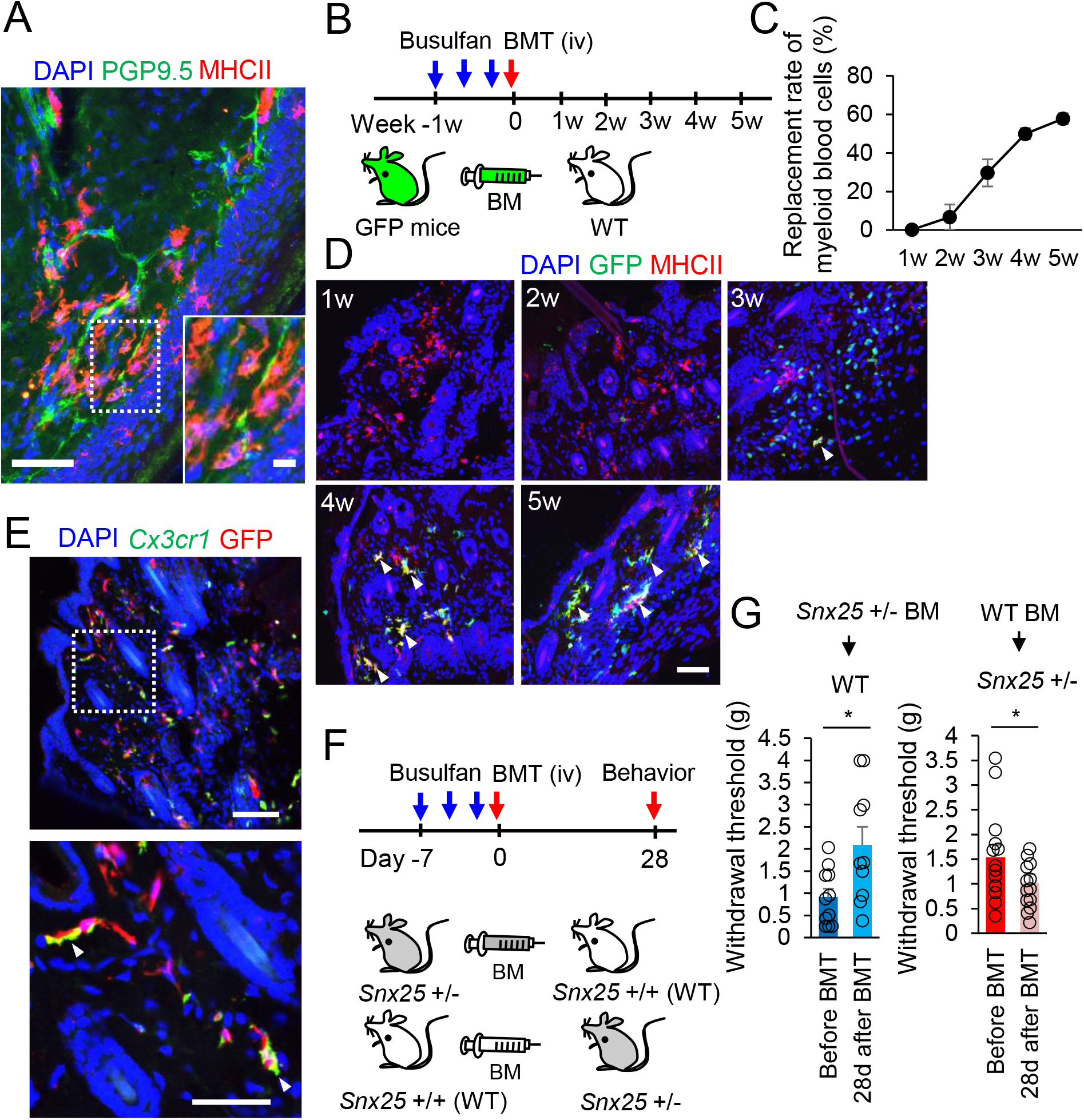
SNX25 in macrophages derived from BM contribute to pain sensation. (**A**) Confocal images of the plantar skin of the hind paw (naive) of WT mice, immunolabeled for PGP9.5 and MHCII. Dermal macrophages (red) are closely associated with PGP9.5-positive nerve (green). Scale bar, 50 μm. The inset at the right lower corner is a magnified view of the boxed area. Scale bar, 10 μm. (**B**) Experimental schedule of bone marrow transplantation (BMT). Bone marrow (BM) chimeric mice were generated by transplanting BM of GFP mice (green mice) to WT mice (intravenous (iv) tail vein injection). (**C**) Chimerism of myeloid cells in peripheral blood, plotted against time after transplantation. n = 3. (**D**) Confocal images of hind paw skin stained for GFP and MHCII in WT mice with BM of green mice. Arrowheads denote double-labeled cells. Double-positive cells increased with time after transplantation. (**E**) Confocal images of hind paw skin stained for GFP (Alexa-594) and *Cx3cr1* mRNA (FISH) in WT mice with BM of green mice (5w after transplantation). BM-derived cells are also positive for *Cx3cr1* mRNA (arrowheads). Scale bar, 100 μm. The lower panel is a magnified view of the boxed area in the upper panel. Scale bar, 50 μm. (**F**) Experimental schedule of cross-transplantation of BMs between WT and *Snx25* +/− mice and behavioral evaluation of chimeric mice. (**G**) Paw withdrawal thresholds to mechanical stimulation with von Frey filaments approached the levels of donors in both chimeras (*Snx25* +/− BM → WT: n = 10; WT BM → *Snx25* +/−: n = 13) compared to thresholds before BMT. Results are represented as mean ± SEM of 3 independent experiments. Significance was calculated using the Student’s *t*-test. \**p* < 0.05.

Unlike tissue-resident macrophages (e.g., microglia in the brain and Kupffer cells in the liver), dermal macrophages in the skin have been shown to be partly derived from bone marrow (BM) and to turn over (Hoeffel et al., 2012), (Tamoutounour et al., 2013), (Kolter et al., 2019). To confirm these features, we transplanted BM from GFP mice (C57BL/6-Tg (CAG-EGFP)) into WT mice using busulfan, which is an efficient reagent to suppress bone marrow cells (Kierdorf et al., 2013). We confirmed that 58% of myeloid blood cells were of donor origin and that donor-derived dermal macrophages expressed MHCII, CD206, and *Cx3cr1* at 5 weeks after BM transplantation (**Figures 3B–3E, Figures S5E** and **S5F**). In contrast, GFP-positive cells were not detected in the gray matter of the spinal dorsal horn, although a few were detected in the pia mater (**Figure S5G**, arrowhead). Consistent with previous reports, these results indicate that spinal cord microglia are not derived from BM in adult (Ginhoux et al., 2010) (**Figure S5G**). To gain further insight into the contribution of dermal macrophages to pain sensation, we made BM chimeric mice by cross-transplanting WT and *Snx25* +/− BMs (**Figure 3F**). Interestingly, the 50% withdrawal threshold to mechanical stimuli in the paws increased in the WT mice with *Snx25* +/− BM transplant and, in turn, decreased in the *Snx25* +/− mice with WT BM transplant (**Figure 3G**). These results strongly suggest that SNX25 in BM-derived immune cells including dermal macrophages, but not spinal microglia, contributes to pain sensation.

The number of macrophages in hind paw skin in *Snx25* +/− mice was normal (**Figure S5B**). However, given that the pain response to a chemical stimulus as an acute inflammatory pain model was extremely reduced in *Snx25* +/− mice (**Figures 2D** and **2E**), we next sought to determine the function of macrophages in the inflammatory environment. Immunohistochemistry revealed that the accumulation of macrophages after formalin injection was reduced in *Snx25* +/− mice (**Figure S6A**). We also found that at 3 d after formalin injection, the expression of a cluster of chemokines was lower in *Snx25* +/− mice than in WT mice (**Figure S6B**). Low macrophage accumulation in *Snx25* +/− mice may be due to this reduction of chemokine expression. These immune phenotypes may be attributable to upregulation of TGF-beta receptor-1 (data not shown), which is known to suppress immune responses (Batlle and Massagué, 2019) and to be degraded by SNX25 (Hao et al., 2011). From these data, we conclude that the abnormality in function of dermal macrophages in *Snx25* +/− mice affects pain sensation under normal and inflammatory conditions.

### *Snx25* conditional KO in macrophages yields a pain-insensitive phenotype

To further target the dermal macrophages, we generated mice having *Snx25* cKO in the monocyte/macrophage lineage by crossing *Snx25^loxP/loxP^* mice with *Cx3cr1*^*Cre*ERT2/WT^ mice (Yona et al., 2013). As expected, *Cx3cr1*^*Cre*ERT2/WT^; *Snx25^loxP/loxP^* mice exhibited reduced pain responses to mechanical and chemical stimuli (**Figures 4A–4C**). Size distribution and the expression of small or large neuron markers in the DRG were normal in *Cx3cr1*^*Cre*ERT2/WT^; *Snx25^loxP/loxP^* mice, indicating that abnormal reactions to pain were not the result of the loss of a particular neuron type (data not shown). However, the mRNA levels of *Scn9a* and *Scn10a* were reduced in the DRG of *Cx3cr1*^*Cre*ERT2/WT^; *Snx25^loxP/loxP^* mice (**Figure 4D**). Consistent with the results in *Snx25* +/− mice, the expression of chemokines and cytokines was lower in hind paw skin of *Cx3cr1*^*Cre*ERT2/WT^; *Snx25^loxP/loxP^* mice than in *Snx25^loxP/loxP^* mice (**Figure 4E**). These results indicate that SNX25 contributes to the inflammatory response in dermal macrophages of skin after chemical stimuli, as well as to pain sensation under normal conditions.

**Figure 4.**
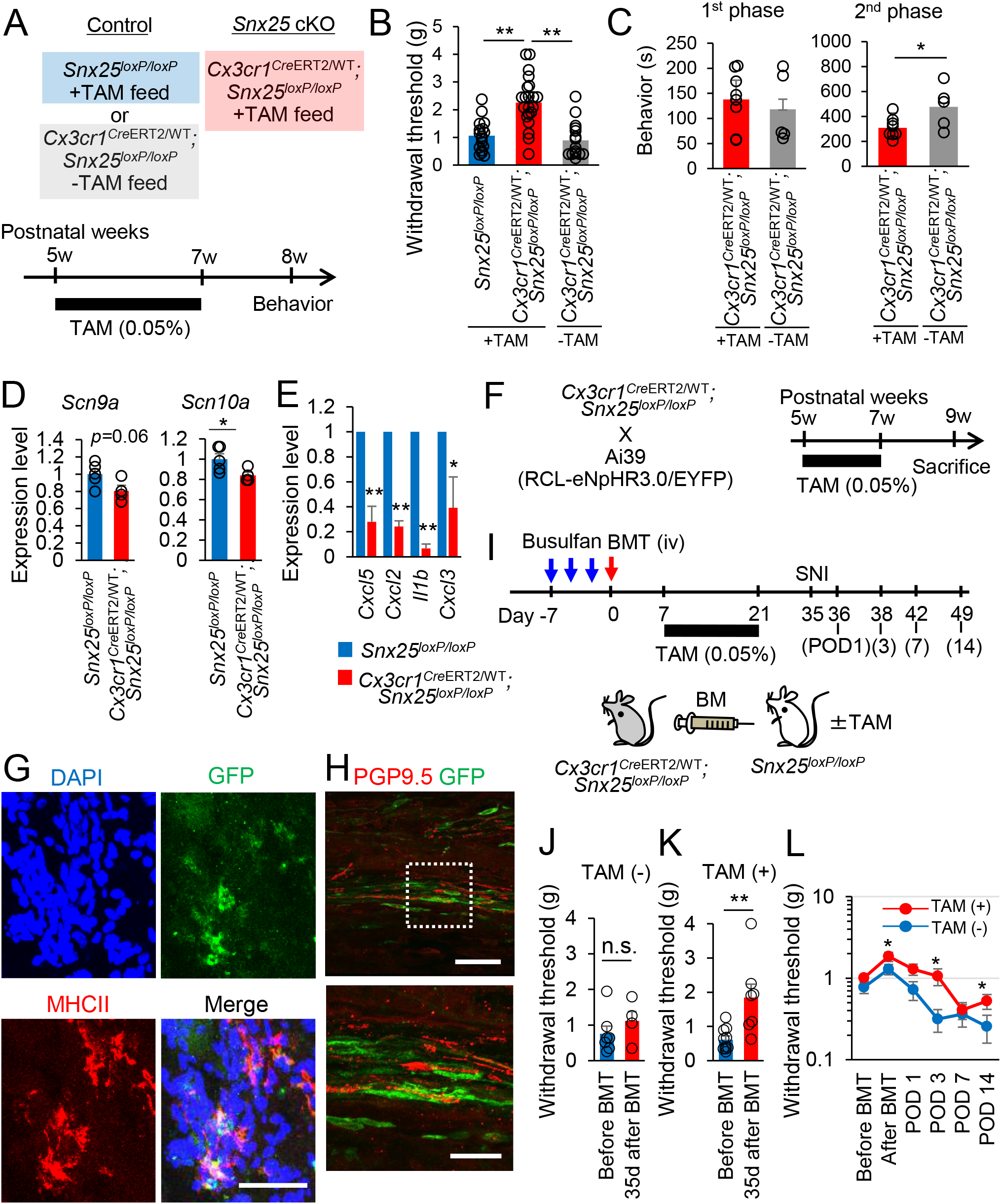
*Snx25* conditional KO in macrophages renders a pain-insensitive phenotype. (**A**) Experimental schedule of conditional KO generation and behavioral analyses. (**B**) von Frey tests demonstrated that *Snx25^loxP/loxP^* mice had pain sensitivity indistinguishable from that of WT mice (Figure 2C). Conditional KO mice (tamoxifen (TAM)-treated *Cx3cr1*^*Cre*ERT2/WT^; *Snx25^loxP/loxP^* mice) exhibited significantly higher thresholds than the controls without tamoxifen treatment or than the floxed mice without the *Cre* driver (*Snx25^loxP/loxP^* mice (+ TAM): n = 17; *Cx3cr1*^*Cre*ERT2/WT^; *Snx25^loxP/loxP^* mice (+ TAM): n = 25; *Cx3cr1*^*Cre*ERT2/WT^; *Snx25^loxP/loxP^* mice (− TAM): n = 14). (**C**) Formalin tests showed that the pain-related behavior time of conditional KO mice was significantly shorter than the control without tamoxifen treatment only in the 2^nd^ phase. (*Cx3cr1*^*Cre*ERT2/WT^; *Snx25^loxP/loxP^* mice (+ TAM): n = 8; *Cx3cr1*^*Cre*ERT2/WT^; S*nx25^loxP/loxP^* mice (− TAM): n = 5). Left, measurement of the behavior time in 1^st^ phase (0–10 min). Right, measurement of the behavior time in 2^nd^ phase (20–60 min). (**D**) Conditional KO DRGs had lower Na channel (*Scn9a* and *Scn10a*) mRNAs than the control DRGs. The difference in *Scn10a* mRNA expression reached statistical significance. (**E**) Chemokines and cytokines were significantly downregulated in the conditional KO skin as compared to the control (*Snx25*-floxed mice without Cre driver). (**F**) Experimental schedule of the visualization of conditional KO cells in the dermis. (**G**) Confocal images of hind paw skin (naive) stained for GFP and MHCII in *Cx3cr1*^*Cre*ERT2/WT^; *Snx25^loxP/loxP^*; Ai39/+ mice. GFP/MHCII double-positive cells (*Snx25* cKO macrophages) are found in the dermis. Scale bar, 50 μm. (**H**) Confocal images of hind paw skin (naive) stained for GFP and PGP9.5 in *Cx3cr1*^*Cre*ERT2/WT^; *Snx25^loxP/loxP^*; Ai39/+ mice. GFP-positive cKO cells are closely associated with the PGP9.5 nerves. Scale bar, 50 μm. The lower panel is a magnified view of the boxed area in the upper panel. Scale bar, 20 μm. (**I**) Schedules for generation of BM chimeric mice by transplanting *Cx3cr1*^*Cre*ERT2/WT^; *Snx25^loxP/loxP^* BM into *Snx25^loxP/loxP^* mice and *Cx3cr1*^*Cre*ERT2/WT^; *Snx25^loxP/loxP^* mice, and subsequent spared nerve injury experiments. (**J**) Even at 35 days after BMT, mechanical pain sensing was comparable to that before BMT if we did not administer tamoxifen (− TAM). (**K**) The same experimental setting as (**J**) except that we treated BM chimeric mice with tamoxifen (+ TAM), yielding a significant increase in withdrawal thresholds to von Frey mechanical stimulation relative to those before BMT. (**L**) Establishment and time course of mechanical allodynia were plotted after BMT and SNI in *Snx25*^*loxP/loxP* (*Cx3cr1-Cre*ERT2/WT; *Snx25*loxP/loxP BM)^ mice with or without tamoxifen treatment (−TAM: n = 4; +TAM: n = 12). Dull sensing was observed at 3 days and 14 days after the SNI operation (postoperative day; POD). Results are represented as mean ± SEM of 3–5 independent experiments. Significance was calculated using one-way ANOVA (B) or the Student’s *t*-test (C, D, J, K, and L) or Welch’s *t*-test (E). \**p* < 0.05, \*\**p* < 0.01.

A recent study demonstrated that Lyve1^lo^MHCII^hi^Cx3cr1^hi^ macrophages colocalize with peripheral nerves (Chakarov et al., 2019). To determine the relationship between SNX25-positive dermal macrophages and peripheral nerves, we crossed *Cx3cr1*^*Cre*ERT2/WT^; *Snx25^loxP/loxP^* mice with reporter mice harboring Rosa-CAG-LSL-eNpHR3.0-EYFP (Ai39) (Madisen et al., 2012) (**Figure 4F**). TAM administration resulted in GFP expression in Cx3cr1/MHCII-positive macrophages (**Figure 4G**), but not in CD117-positive mast cells (data not shown). GFP-positive dermal macrophages were apposed to PGP9.5-positive fibers in the dermis (**Figure 4H**), suggesting that SNX25-positive dermal macrophages are associated with peripheral sensory fibers.

CX3CR1 is the fractalkine receptor and is found not only on the surface of macrophages but also on the surface of microglia in the central nervous system (Fumagalli et al., 2013). Microglia also regulate neuronal and synaptic activities to change pain behavior (Tsuda et al., 2003), implying that the pain-insensitive phenotype in the *Cx3cr1*^*Cre*ERT2/WT^; *Snx25^loxP/loxP^* mice is derived from a microglial abnormality rather than from dermal macrophage dysfunction. To distinguish between the cells responsible (dermal macrophages or microglia) in the *Cx3cr1-Cre* driven *Snx25* cKO mice, we again transplanted BM of *Cx3cr1*^*Cre*ERT2/WT^; *Snx25^loxP/loxP^* mice into *Snx25^loxP/loxP^* mice (**Figure 4I**). In these BM chimeric mice, *Snx25* cKO was limited to dermal macrophages; we confirmed that BM-derived cells did not contribute to the microglia in the spinal cord (**Figure S5G**). This finding is consistent with an *in vivo* lineage-tracing study demonstrating that adult microglia are derived from primitive myeloid progenitors before embryonic day 8 (Ginhoux et al., 2010). Notably, the withdrawal threshold to mechanical stimuli was significantly increased in the *Snx25^loxP/loxP^* mice having *Cx3cr1*^*Cre*ERT2/WT^; *Snx25^loxP/loxP^* mice BM transplanted and TAM administered at 35 d after transplantation (**Figure 4K**), while the same experimental condition without TAM treatment yielded a threshold comparable to the control (before BM transplantation, **Figure 4J**). Furthermore, in spared nerve injury (SNI) paradigms using the same experimental (+TAM) and control (−TAM) mice as above, TAM treatment attenuated mechanical hypersensitivities that were observed in the control group (**Figure 4L**). These results indicate that SNX25 in dermal macrophages, but not microglia, is required for pain sensation under both normal and painful conditions.

### SNX25 in dermal macrophages is required for pain sensation via NGF signaling

We focused on nerve growth factor (NGF) as a critical factor in the pain-insensitive behavior of mice having *Snx25* cKO in dermal macrophages for two reasons: first, NGF plays critical roles in hyperalgesia and its mutation causes painless phenotypes (Hefti et al., 2006), (Mantyh et al., 2011); and second, *Trpv1*, *Scn9a*, and *Scn10a*, whose expression was reduced in the *Snx25* +/− DRGs (**Figure 2M**), are all transcriptionally regulated by peripheral tissue-derived and retrogradely transported NGF (Mantyh et al., 2011). A plausible scenario is that NGF concentration in the dermis is partly maintained by macrophages in WT mice and that decreased NGF impairs macrophage-to-nerve signaling in the *Snx25* heterozygotes and cKO in macrophages. Consistent with this, the NGF expression level of hind paw skin was decreased in the *Snx25* +/− mice (**Figure 5A**). We also found that the expression level of NGF was lower at 30 min after formalin injection in hind paw skin of *Snx25* +/− mice (**Figure 5B**). NGF was actually expressed in dermal macrophages in hind paw skin (**Figure 5C**) and its expression level was reduced in BM-derived macrophages (BMDMs) of *Snx25* +/− mice (**Figure 5D**). We further conducted a nerve ligation assay to assess the cumulative axonal transport rate of TrkA, which is the cognate receptor for NGF, in sciatic nerves (Chaumette et al., 2020). Eight hours after ligation, perfused sciatic nerve tissues were analyzed immunohistochemically. The accumulation of TrkA receptor on the distal side of the nerve ligature was significantly reduced in the *Snx25* +/− nerves (**Figure 5E**), supporting the notion that a reduction of NGF in the periphery results in diminished retrograde transportation of the NGF–TrkA complex in the *Snx25* +/− DRG. Quantitative RT-PCR showed decreased *Ngf* mRNA in BMDMs from *Snx25* +/− mice or in BMDMs with *Snx25* knockdown (KD), indicating that SNX25 modulates *Ngf* production at the mRNA level (**Figures 5F** and **5G**).

**Figure 5.**
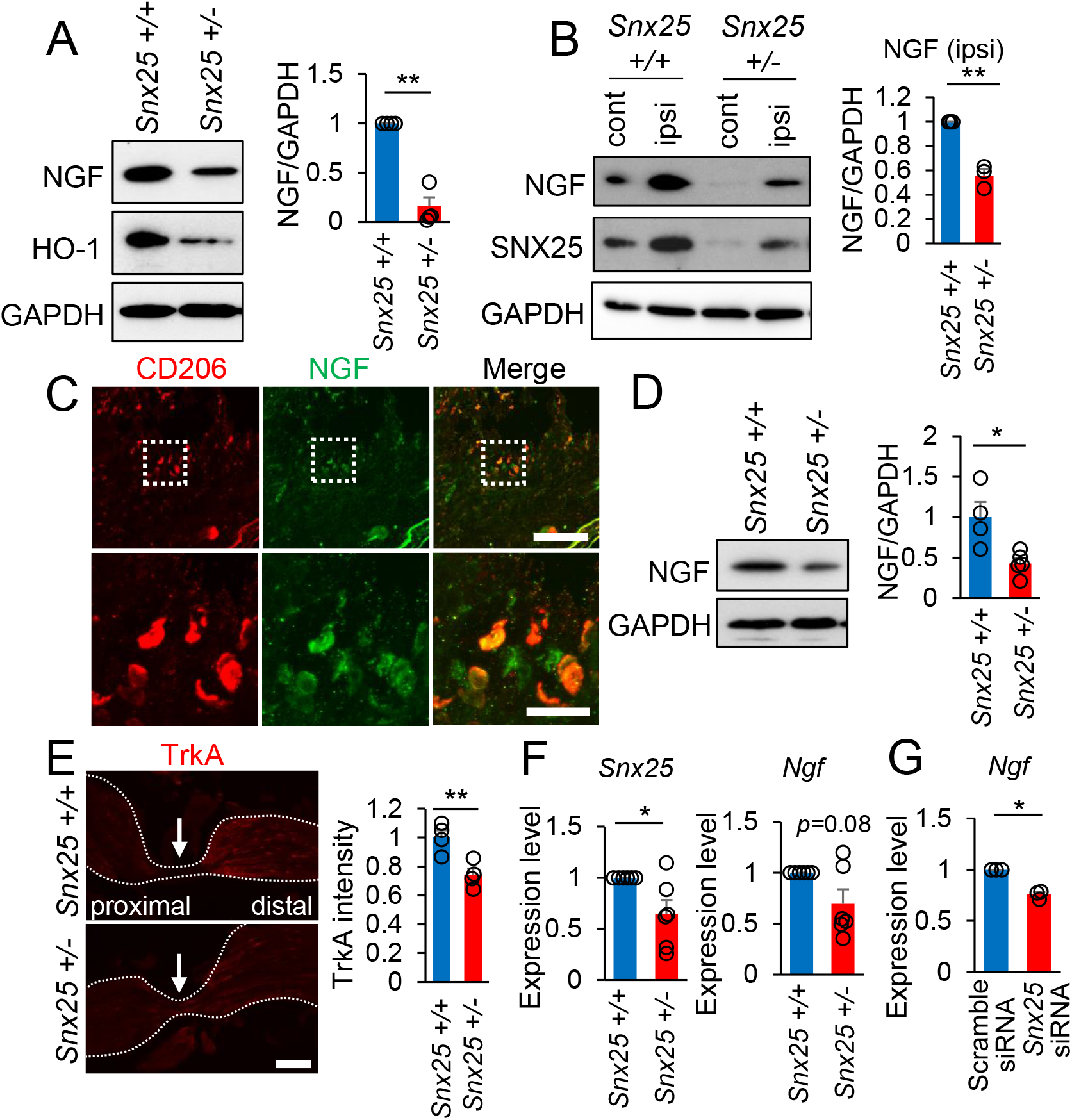
NGF expression in macrophages is reduced in *Snx25* +/− mice. (**A**) A representative Western blot showing NGF levels in the hind paw skin of WT and *Snx25* +/− mice. Heme oxigenase-1 (HO-1), a stress-inducible and downstream gene of Nrf2 transcription factor, was probed on the same blot as the positive control. The graph shows semi-quantitative analyses of Western experiments. NGF content normalized with GAPDH expression was significantly lower in the heterozygote than in the WT mice. (**B**) Semi-quantitative analyses of NGF levels in the hind paw skin of WT and *Snx25* +/− mice at 30 min after formalin injection. Both NGF and SNX25 were increased by formalin injection in both WT and heterozygote mice, but the protein levels were lower in the *Snx25* +/− mice when we compared the same sides. The graph shows that NGF levels in the ipsilateral side (formalin-injected side) were significantly decreased in the heterozygote mice. (**C**) Confocal images of hind paw skin immunolabeled for NGF and CD206 in WT mice. Scale bar, 200 μm. Lower panels are magnified views of the boxed areas in the upper panels. Scale bar, 50 μm. Note the clear colocalization of CD206 and NGF. (**D**) Expression levels of NGF in BMDMs of WT and *Snx25* +/− mice were examined by Western blotting, normalized with GAPDH content, and analyzed semi-quantitatively (WT: n = 4; *Snx25* +/−: n = 5). (**E**) Left, confocal images of sciatic nerve sections immunolabeled for TrkA at 8 h after nerve ligation (arrows indicate ligation site) in WT and *Snx25* +/− mice. Right, semi-quantitative analysis of the TrkA accumulation on the distal side of the nerve ligature. Note the significant decrease of TrkA protein in the heterozygote mice. Scale bar, 200 μm. (**F**) Expression profiles of mRNAs for *Snx25* and *Ngf* in BMDMs of WT and *Snx25* +/− mice (WT: n = 6; *Snx25* +/−: n = 6). (**G**) *Ngf* mRNA was quantified by RT-PCR in BMDMs transfected with either *Snx25* siRNA or scramble siRNA (scramble siRNA: n = 3; *Snx25* siRNA: n = 3). *Snx25* knockdown led to a significant decrease in *Ngf* mRNA expression. Results are represented as mean ± SEM of 3–5 independent experiments. Statistical analyses were performed using the Student’s *t*-test (D and E) or Welch’s *t*-test (A, B, F and G). \**p* < 0.05, \*\**p* < 0.01.

We next investigated the molecular mechanisms bridging SNX25 to *Ngf* synthesis. A CNC-bZip transcription factor, NF-E2-related factor 2 (Nrf2), regulates *Ngf* mRNA induction in glial cells (Mimura et al., 2011). Consistent with that report, we found that *Nrf2*-specific siRNA significantly reduced constitutive *Ngf* gene expression in BMDMs (**Figure 6A**). We hypothesized that SNX25 regulates Nrf2 level and thereby *Ngf* gene expression. Nrf2 level in the cell is known to be regulated by continuous ubiquitination and proteasome degradation, which is blocked by Keap1 protein (Kensler et al., 2007). The level of poly-ubiquitinated Nrf2 protein was increased following treatment with a proteasome inhibitor, MG132 (**Figure 6B**, arrowhead) and was further elevated by siRNA-mediated knockdown of *Snx25* in 293T cells (**Figure 6C**, arrowheads). *Snx25* overexpression (cells transiently co-transfected with mouse *Snx25* and mouse *Nrf2* expression vectors), in turn, decreased poly-ubiquitinated Nrf2 level compared to empty vector (cells transiently co-transfected with empty vector and mouse *Nrf2* expression vector) in 293T cells (**Figure 6D**, arrowhead). The soluble-type tamoxifen derivative 4-OH tamoxifen (4-OHT) conditionally decreased *Snx25* in *Cx3cr1*^*Cre*ERT2/WT^; *Snx25^loxP/loxP^* BMDMs, and this *in vitro* cKO recapitulated the above-demonstrated increase of poly-ubiquitinated Nrf2 (**Figure 6E**, arrowhead). These results indicate that SNX25 activates NGF production by inhibiting ubiquitin-mediated degradation of Nrf2. In support of this interpretation, a representative target factor of Nrf2, heme oxygenase-1 (HO-1), was decreased in hind paw skin of the *Snx25* +/− mice (**Figure 5A**).

**Figure 6.**
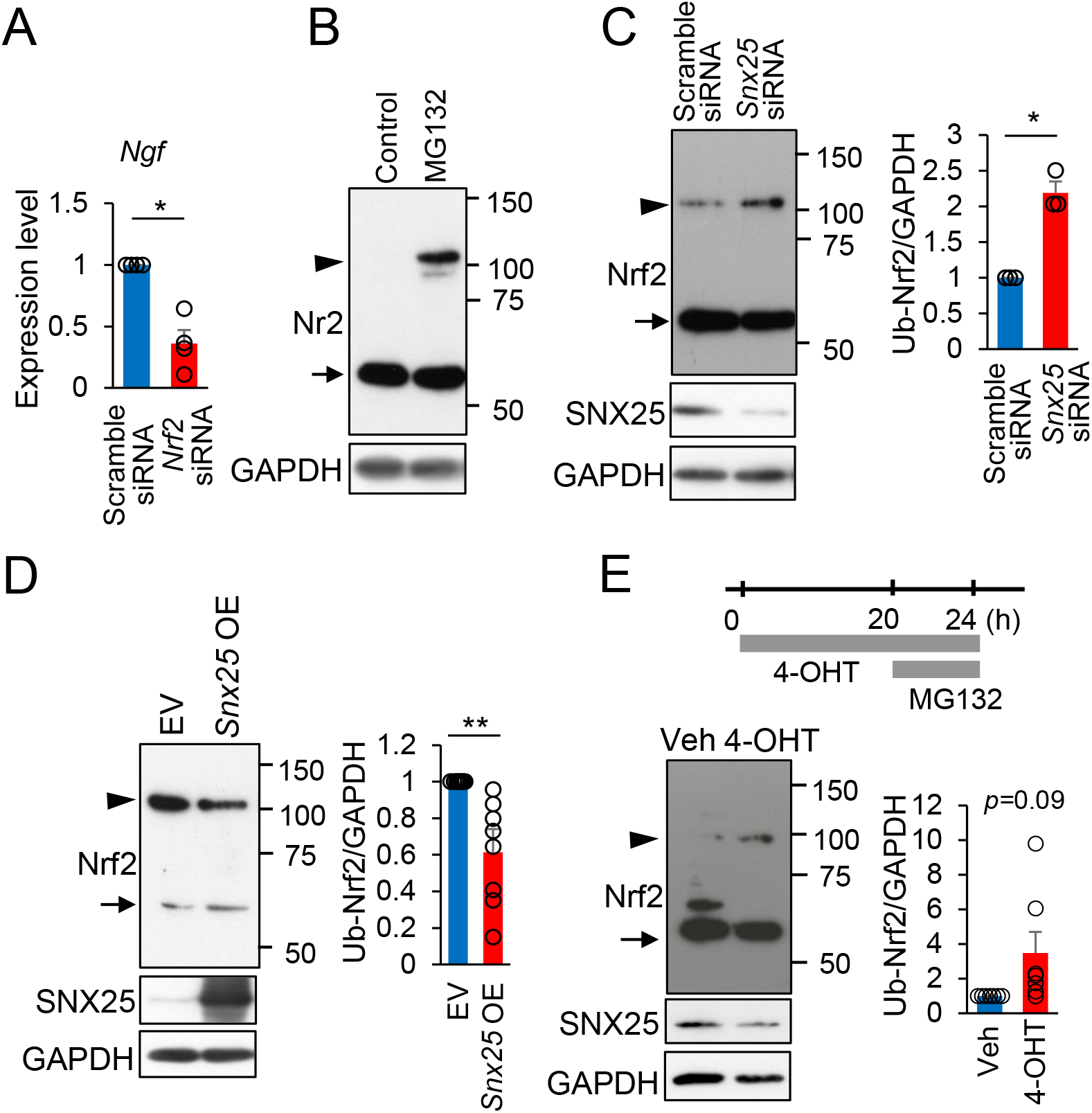
SNX25 activates *Ngf* production by inhibiting ubiquitin-mediated degradation of Nrf2. (**A**) *Ngf* mRNA levels in BMDMs transfected with either *Nrf2* siRNA or scramble siRNA were analyzed semi-quantitatively by RT-PCR. *Nrf2* knockdown resulted in a significant decrease of *Ngf* mRNA (scramble siRNA: n = 4; *Snx25* siRNA: n = 4). (**B**) A representative Western blot showing Nrf2 protein levels in BMDMs in the presence or absence of MG132 (5 μM, 4 h). Arrow: Nrf2 (61–68 kDa); arrowhead: poly-ubiquitinated Nrf2 (100–110 kDa). (**C**) Ubiquitination levels of Nrf2 protein were probed in 293T cells transfected with *Snx25* siRNA or scramble siRNA in the presence of MG132 (5 μM, 4 h) (scramble siRNA: n = 3; *Snx25* siRNA: n = 3). Arrow: Nrf2 (61-68 kDa); arrowhead: poly-ubiquitinated Nrf2 (100–110 kDa). Right, semi-quantitative analysis revealed that the band intensity of poly-ubiquitinated Nrf2 (Ub-Nrf2) is significantly higher in the *Snx25* knock-down cells than in the control (scramble siRNA-transfected). (**D**) Nrf2 and poly-ubiquitinated Nrf2 were probed in 293T transfected with full-length *Snx25* expression vector (*Snx25* OE) or empty vector (EV) in the presence of MG132 (5 μM, 4 h) (empty vector + *Nrf2* vector: n = 8; *Snx25* vector + *Nrf2* vector: n = 8). Arrow: Nrf2 (61-68 kDa); arrowhead: poly-ubiquitinated Nrf2 (100–110 kDa). Right, Semi-quantitative analysis shows that the intensity of poly-ubiquitinated Nrf2 (Ub-Nrf2) in the *Snx25*-overexpressing cells was significantly lower than that in the cells transfected with empty vector. (**E**) Upper panel shows the experimental schedule. Lower left panel: Nrf2 (non-ubiquitinated and ubiquitinated species) was examined by Western analyses in BMDMs of *Cx3cr1*^*Cre*ERT2/WT^; *Snx25^loxP/loxP^* treated with 4-OH-tamoxifen (4-OHT, 1 μM, 24 h) or without 4-OHT (vehicle (Veh)) in the presence of MG132 (5 μM, 4 h) (Veh: n = 7; 4-OHT: n = 7). Arrow: Nrf2 (61-68 kDa); arrowhead: poly-ubiquitinated Nrf2 (100–110 kDa). Lower right panel: quantitative analysis of the intensity of poly-ubiquitinated Nrf2 (Ub-Nrf2). 4-OHT treatment significantly increased ubiquitinated species of Nrf2. Results are represented as mean ± SEM of 3–5 independent experiments. Statistical analyses were performed using the Welch’s *t*-test. \**p* < 0.05, \*\**p* < 0.01.

An important question is whether dermal macrophages are sufficient to initiate pain sensation without both neuropathic intervention and inflammation. To address this, we depleted dermal macrophages by intradermal injection (i.d.) of clodronate liposomes (Clo-lipo), a well-characterized macrophage killer (Ding et al., 2019), twice into one side of the hind paw (**Figures 7A** and **B**). Immunohistochemistry revealed that the numbers of CD206- or MHCII-positive macrophages were decreased at 3 days after the second Clo-lipo injection relative to the control liposome-injected skin (**Figure 7C**). Notably, macrophage depletion increased withdrawal thresholds to mechanical stimuli (**Figure 7E**), while control liposomes did not (**Figure 7D**). Western blot analyses revealed that NGF, SNX25, and CD206 expression levels were decreased lower on the Clo-lipo-injected area than on the control liposome-injected area (**Figures 7F–I**). Taken together, these findings indicate that SNX25 and NGF in dermal macrophages are required for pain sensation under normal conditions.

**Figure 7.**
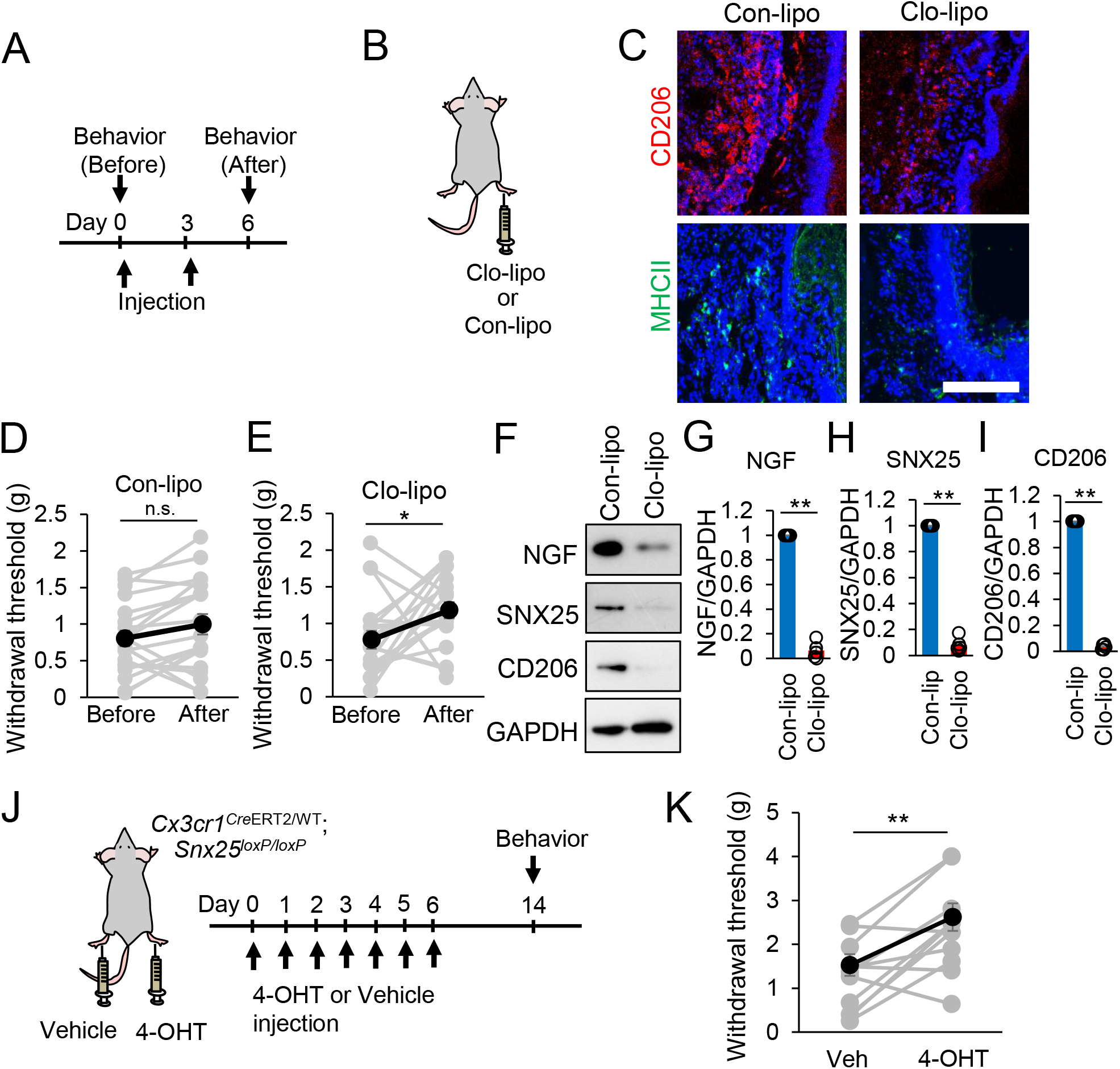
Dermal macrophages are sufficient to initiate pain sensation and SNX25 is a key factor in this process. (**A**) Experimental schedule of macrophage killing in the dermis. von Frey tests were conducted before and after the liposome injections. (**B**) Control liposomes (Con-lipo) and clodronate liposomes (Clo-lipo) were injected into hind paws. An examiner who was blind to the side of injection assessed mechanical pain sensitivity with the von Frey test. (**C**) Confocal images of hind paw skin immunolabeled for CD206 (red) or MHCII (green) in WT mice injected with Con-lipo or Clo-lipo. Note that CD206-positive or MHCII-positive dermal macrophages were decreased in the Clo-lipo-injected skin. Scale bar, 200 μm. (**D**) Paw withdrawal thresholds to von Frey filaments on the Con-lipo-injected side (n = 20) were at comparable levels before and after injection. (**E**) Paw withdrawal thresholds to von Frey filaments on the Clo-lipo-injected side (n = 20) were significantly elevated as compared to those before injection. (**F–H**) Expression levels of NGF and SNX25 in the hind paw skin of the Con-lipo-injected and Clo-lipo-injected sides were examined by Western blotting (**F**) and semi-quantitatively compared for NGF (**G**) and SNX25 (**H**) and CD206 (**I**). Both NGF and SNX25 were decreased on the Clo-lipo-injected sides (n = 5). (**J**) Scheme depicting dermal injection of vehicle or 4-OH-tamoxifen (4-OHT) into hind paws of a *Cx3cr1*^*Cre*ERT2/WT^; *Snx25^loxP/loxP^* mouse and experimental time course. To achieve gene recombination, we injected 4-OHT daily for 7 days and pain behavior was evaluated at 1 week after last injection. (**K**) Paw withdrawal thresholds to von Frey filaments are compared between the vehicle- and 4-OHT-injected sides (n = 11). The 4-OHT-injected side was insensitive to mechanical stimulation as compared to the vehicle-injected side. Results are represented as mean ± SEM of 3 independent experiments. Statistical analyses were performed using the Student’s *t*-test (D, E and K) or Welch’s *t*-test (G, H and I). \**p* < 0.05, \*\**p* < 0.01.

To further substantiate the importance of SNX25 in pain sensation, we administered 4-OHT (40 ng/μL, 10μL) by intradermal injection daily for seven days into *Cx3cr1*^*Cre*ERT2/WT^; *Snx25^loxP/loxP^* mice. Vehicle was injected into the contralateral side of the same animal (**Figure 7J**). At 8 days after the last injection, the 4-OHT-injected hind paw showed a pain-insensitive phenotype, in contrast to the vehicle-injected hind paw (**Figure 7K**). These data indicate that SNX25 in dermal macrophages is a pivotal factor for pain sensation under normal conditions.

Collectively, our results indicate that SNX25 activates NGF production by inhibiting ubiquitin-mediated degradation of Nrf2, resulting in increased expression of a number of pain-related genes in the DRG cell bodies (**Figure S7**). Based on these data, we conclude that SNX25 in dermal macrophages modulates acute pain sensing under normal and painful conditions.

## Discussion

Pain significantly reduces quality of life in various diseases. Recent pain research has revealed pain-modulating cells, molecules, and neural pathways, especially in the central nervous system (Grace et al., 2014) (Ji et al., 2016) (Inoue and Tsuda, 2018) (Matsuda et al., 2019). The functional anatomy (Crawford and Caterina, 2020) and neuro-immune interactions (Ren and Dubner, 2010) (Malcangio, 2019) of the peripheral sensory system have been elucidated at remarkable speed in recent years. However, the whole picture of pain-sensing mechanisms still remains unclear. In the present study, from a phenotype-driven forward genetic screen of pain-insensitive *Mlc1* TG mice that was free from any specific working hypothesis, we have successfully identified *Snx25* as a pain-modulating gene. Both *Snx25* +/− mice and *Snx25* conditional-KO mice in macrophages displayed reduced pain responses under both normal and painful conditions (**Figures 2 and 4**). SNX25 inhibits the ubiquitination and subsequent proteasome degradation of Nrf2 and thereby maintains NGF production and secretion into tissues. *Snx25* conditional KO, in turn, accelerates Nrf2 degradation and lowers NGF levels, which leads to a dull phenotype (**Figure S7**).

Recent progress in gene cataloging techniques such as single-cell RNA sequencing has broadened our knowledge of tissue macrophages. Chakarov et al. characterized two independent populations of lung interstitial macrophages exhibiting distinct gene expression profiles and phenotypes: Lyve1^lo^MHCII^hi^Cx3cr1^hi^ macrophages were associated with nerves, whereas Lyve1^hi^MHCII^lo^Cx3cr1^lo^ macrophages were preferentially located around blood vessels (Chakarov et al., 2019). These interstitial macrophages were in part derived from bone marrow (Chalarov et al., 2019), consistent with fate-mapping studies (Tamoutounour et al., 2013) (Kolter et al., 2019). We confirmed that donor-derived GFP-positive cells also expressed *Cx3cr1* and MHCII in the dermis of recipient mice after BM transplantation (**Figures 3B–3E**). We crossed *Cx3cr1*^*Cre*ERT2/WT^ mice with the reporter line, Rosa-CAG-LSL-eNpHR3.0-EYFP (Ai39) (Madisen et al., 2012), and GFP-positive dermal macrophages were frequently found in close proximity to PGP9.5-positive fibers that innervate the skin (**Figure 4H**). This finding is consistent with a previous report that Cx3cr1^hi^ macrophages colocalize with peripheral nerves, which contributes to the surveillance and regeneration of local nerves in the dermis (Kolter et al., 2019). NGF production by these dermal macrophages (**Figure 5C**) likely contributes to the regeneration of local nerves in addition to the maintenance of pain sensibility. Macrophages are known to adhere to the cell matrix at a specialized structure, the podosome (van den Dries et al., 2019). Podosomes confer on dermal macrophages the ability to sense mechanical stress and deformation of tissues (Alonso et al., 2019) (van den Dries et al., 2019). It is interesting to speculate that the mechanosensing ability of dermal macrophages is linked to NGF production and thereby regulates mechanical pain sensitivity. In this context, we observed a tendency for SNX25 and NGF to be upregulated in response to mechanical stretch in cultured bone marrow-derived macrophages (data not shown).

We showed that SNX25 regulated cellular Nrf2 content by changing its ubiquitination level (**Figure 6**). Although SNX family members are diverse and are involved in a wide variety of intracellular events such as regulation of vesicle trafficking (Teasdale and Collins, 2012), some of them regulate protein ubiquitination. SNX16 inhibits ubiquitin-mediated proteasomal degradation of eukaryotic translation elongation factor 1A2 in colorectal cancer development (Shen et al., 2020). SNX17 recruits USP9X to antagonize ubiquitination and degradation of pericentriolar material 1 during serum starvation-induced ciliogenesis (Wang et al., 2019). We also found that *Snx25* knockdown in the macrophage cell line RAW264.7 promoted ubiquitination of IκBα after lipopolysaccharide stimulation (unpublished data). Nrf2 is the principal transcription factor that regulates antioxidant response element-mediated expression of antioxidant enzymes (Kensler et al., 2007). Recent studies have reported a relationship between Nrf2 and mechanical stimuli (Rysä et al., 2018) (Li et al., 2018). Taking all these observations together, mechanosensory stimuli impinging on skin may stimulate dermal macrophages, and the macrophages then make the NGF concentration optimal for neurons to respond to stimuli via an SNX25–Nrf2 signaling pathway.

One of the most important findings in the present study is that lowering NGF levels for a relatively short term (within one month) yielded mice with the pain-insensitive phenotype in *Snx25* cKO in dermal macrophages (**Figures 7J** and **7K**) and in mice transplanted with *Snx25* cKO macrophages (**Figure 4I-L**) in naïve glabrous skin. In human, hereditary sensory and autonomic neuropathy type V (HSAN V), characterized by a marked absence of pain sensibility, is caused by mutations in the *Ngf* gene (Einarsdottir et al., 2004) (Capsoni, 2014). The NGF suppression by *Snx25* cKO in macrophages mimics HSAN V pathology to some extent, but there is a critical difference between two paradigms: HSAN V is characterized by long-term NGF deficiency and morphological changes in peripheral sensory nerves, such as retraction of nerve endings (Axelsson et al., 2009), which we did not see in the *Snx25* cKO dermis. A short-term NGF decrease in *Snx25* cKO mice depletes the pain-sensing machinery (Na channels) in DRG neurons (**Figures 4D and 4E**). Based on the clinical phenotypes of HSAN V patients, anti-NGF neutralizing monoclonal antibodies were developed as therapeutic means to mitigate refractory pain (Wild et al., 2007) (Röhn et al., 2011) (Zhou et al., 2019). Humanized monoclonal antibodies (tanezumab and fasinumab) have gone to clinical trials with successful pain-relieving effects (Bannwarth and Kostine, 2014) (Hefti, 2020). Although an unexpected side effect on joints precluded the monoclonal antibodies to further proceed to bedside, they are still a good target of pain-relieving medicine (Hefti, 2020). The increase of tissue NGF levels, on the other hand, is well characterized in several inflammatory conditions and in several models of pain (Woolf et al., 1994). Indeed, we showed that the expression level of NGF was elevated at 30 min after formalin injection in hind paw skin (**Figure 5B**). Taking all these findings into account, we propose that the tissue (dermis in the present study) content of NGF is continuously controlled at least in part by macrophages through SNX25–Nrf2 signaling. In this hypothesis, the tissue levels of NGF parallel the mechanical pain sensitivities: the higher the NGF level, the more sensitive the animal or tissue is, and vice versa. Supporting this hypothesis, a Clo-lipo-mediated purge of dermal macrophages led to lowered NGF levels and concomitant pain-insensitive phenotypes (**Figures 7F–7I**). SNX25–Nrf2 signaling-mediated NGF regulation broadens the role of dermal macrophages. The relationship between macrophages and pain sensation has long been examined and most studies have focused on pathological painful situations; for example, complement C5a stimulates macrophages and thereby causes mechanical hypersensitivity (Warwick et al., 2019) or thermal hypersensitivity (Shutov et al., 2016), both of which are inflammatory conditions. Angiotensin also causes hyperalgesia through macrophage stimulation (Shepherd et al., 2018a) (Shepherd et al., 2018b). These studies did not check the pain sensitivity in naïve conditions. SNX25–Nrf2 signaling in macrophages has the potential to bridge between the painless phenotype of HSAN V and these hyperalgesia conditions. In due course, it would be tempting to develop small compound(s) that could inhibit the SNX25–Nrf2 signaling pathway. Such compounds may become a promising alternative to the anti-NGF monoclonal antibodies, which have been withheld from clinical applications. We should, however, exercise caution in taking this step; NGF is also produced by noninflammatory cells, such as keratinocytes (Tron et al., 1990) and endothelial cells (Foster et al., 2003), in addition to other inflammatory cells, such as fibroblasts. Thus, further experiments are needed to determine the entire cellular and molecular mechanism controlling peripheral NGF levels.

## Acknowledgements

We thank Kazunori Sango (Tokyo Metropolitan Institute of Medical Science), Shenglan Wang (Hyogo University of Health Sciences), Toshihiro Ito (Nara Medical University), Masahiro Kitabatake (Nara Medical University), Kazuki Nakahara (Nara Medical University), and Yoshie Kawabe (Nara Medical University) for technical assistance. This work was supported by JSPS KAKENHI JP16K08451 (to HO), JP16K20112 (to YT), JP18K16492 (to TS), JP19K07827 (to TT), JP19K18303 (to YT), JP19K16480 (to AI), the Osaka Medical Research Foundation for Intractable Diseases (to TT), the Takeda Science Foundation (to TT), the Nakatomi Foundation (to TT), and the Naito Foundation (to TT).

## Author contributions

TT, HO, HF, and AW conceived the project and designed the experiments. TT, HO, YT, TS, MB, ST, KN, AI, ST, and KT performed the experiments. TT, HO, MB, and KT analyzed the data. TT and AW wrote the paper. AW coordinated and directed the project.

## STAR Methods

### Animals

*Mlc1* TG mice (B6; CBB6(129)-Tg(Mlc1-tTA)2Rhn) were a gift from K.F. Tanaka (Keio University). *SNX25* constitutive KO (*Snx25* +/−) mice (B6/N-*Snx25*^tm1a/Nju^, Strain number T001400) were obtained from Nanjing BioMedical Research Institute of Nanjing University (NBRI). *Snx25* cKO mice were generated by first crossing our *Snx25* LacZ/+ mice with CAG-Flpo mice (B6.Cg-Tg(CAG-FLPo)/1Osb), which were a gift from M. Ikawa (Osaka University), in order to excise the LacZ cassette framed by *Frt* sites and obtain an allele with floxed exon 4 (*Snx25^loxP/loxP^* mice) (Yamazaki et al., 2016). We crossed *Advillin*-cre mice (B6.Cg-Tg(*Avil*-*Cre*/ERT2)AJwo/J) (Jackson Laboratory, Stock No: 032027) with *Snx25^loxP/loxP^* mice to obtain *Avil^Cre^*^ERT2/WT^; *Snx25^loxP/loxP^* mice. We crossed *Cx3cr1*^*Cre*ERT2^ mice (B6.129P2(C)-*Cx3cr1*^tm2.1(*Cre*/ERT2)Jung/J^) (Jackson Laboratory, Stock No: 020940) with *Snx25^loxP/loxP^* mice to obtain *Cx3cr1*^*Cre*ERT2/WT^; *Snx25^loxP/loxP^* mice. We crossed *Cx3cr1*^*Cre*ERT2/WT^; *Snx25^loxP/loxP^* mice with reporter mice Rosa-CAG-LSL-eNpHR3.0-EYFP (Ai39 mice, Jackson Laboratory, Stock No: 014539) to obtain *Cx3cr1*^*Cre*ERT2/WT^; *Snx25^loxP/loxP^*; Ai39 mice. C57BL/6-Tg (CAG-EGFP) mice were purchased from Japan SLC (Hamamatsu, Japan). They were housed in standard cages under a 12 h light/dark cycle and temperature-controlled conditions. All the protocols for the animal experiments were approved by the Animal Care Committee of Nara Medical University in accordance with the policies established in the NIH Guide for the Care and Use of Laboratory Animals. This study was also carried out in compliance with the ARRIVE guidelines (https://arriveguidelines.org/).

### Behavioral test

Paw mechanical sensitivity was assessed using von Frey’s filaments based on the up-down method developed by Chaplan (Chaplan et al., 1994). The von Frey’s filaments used were: 0.07, 0.16, 0.4, 0.6, 1, 1.4, 2, 4 g. Animals were acclimatized for at least 15 min in individual clear acrylic cubicles (10 × 10 × 10 cm) placed on top of an elevated wire mesh. Quick withdrawal or licking of the paw after the stimulus was considered a positive response. Threshold values were derived according to the method described by Chaplan (Chaplan et al., 1994). For the formalin test, 10☐μl of 5% formalin was injected subcutaneously into the plantar surface of the right hind paw. PBS (10☐μl) was injected into the plantar surface of the left hind paw. We calculated the durations of lifting, shaking, and licking of the formalin-injected paw. For hot plate test, mice were acclimatized for at least 2 h (1h/day x 2days) in individual clear acrylic cubicles placed on the preheated plate. The withdrawal latency in response to the stimulus was determined manually. In all the behavioral tests, examiners were always blind to the genotypes of mice, the kinds of treatments, and the sides of hind paws that received injections. After the evaluation was done, the behavioral data were analyzed by a different researcher.

### Surgery of spared nerve injury (SNI) model

The surgery of the SNI model was conducted as described previously (Decosterd and Woolf, 2000). Surgical procedures were performed under 2% isoflurane anesthesia. SNI was made by a 6-0 polypropylene thread with tight ligation of the two branches of the right sciatic nerve, the common peroneal and the tibial nerves, followed by transection and removal of a 2-mm nerve portion. The sural nerve remained intact and any contact with or stretching of this nerve was carefully avoided. Muscle and skin were closed in two distinct layers.

### Reagents

For tamoxifen (TAM) treatment, we employed oral administration. TAM (Sigma-Aldrich, St. Louis, MO, USA) was mixed with powdered chow (0.5 mg/g normal chow). This oral administration method is convenient for continuous administration and results in efficient induction of recombination while minimizing stress on the mice (Kiermayer et al., 2007). For *Snx25* deletion in BMDMs, we treated cells with *Snx25*-specific siRNA (Sigma Aldrich) using Lipofectamine RNAiMAX transfection reagent (Thermo Fisher). For *Snx25* deletion in BMDMs derived from *Cx3cr1*^*Cre*ERT2/WT^; *Snx25^loxP/loxP^* mice, we treated cells with 1 μM 4-OH-tamoxifen (4OHT, Sigma Aldrich) for 24 h. For inhibition of proteasomes, we used 5 μM MG132 (Sigma Aldrich) for 4 h.

### Immunohistochemistry

Sections were immersed in PBS containing 5% bovine serum albumin and 0.3% Triton X-100 for 1 h. Antibodies against mouse anti-CGRP (1:500, ab1887, abcam, Cambridge, UK), rabbit anti-c-Fos (1:10000, 226003, Synaptic Systems, Gottingen, Germany), rabbit anti-TRPV1 (1:100, KM018, Trans Genic, Fukuoka, Japan), rabbit anti-TrkA (1:150, ab76291, abcam), mouse anti-NF200 (1:1000, N0142, Sigma-Aldrich), mouse anti-PGP9.5 (1:500, ab8189, abcam), rat anti-MHCII (1:100, NBP1-43312, Novus Biologicals, Centennial, CO, USA), goat anti-CD206 (1:500, AF2535, R&D Systems, Minneapolis, MN, USA), rabbit anti-Iba1 (1:500, 019-19741, Wako, Osaka, Japan), rabbit anti-NGF (1:1000, sc-548, Santa Cruz Biotechnology, Santa Cruz, CA, USA), rat anti-GFP (1:5000, 04404-84, nacalai tesque, Kyoto, Japan), rabbit anti-GFP (1:5000, A6455, Thermo Fisher Scientific, Waltham, MA, USA), rabbit anti-SNX25 (1:500, 13294-1-AP, Proteintech, Rosemont, IL, USA) were applied overnight at 4°C. Alexa Fluor 488- and 594- (1:1000, Life Technologies, Grand Island, NY, USA) conjugated IgG were used as secondary antibodies. Sections were subjected to fluorescent Nissl staining (Neurotrace, Molecular Probes, Eugene, OR, USA). Images were captured using a confocal laser scanning microscope (C2, Nikon, Tokyo, Japan).

### Microarray

Total RNA was isolated from the bone marrow of C57BL/6 mice and *Mlc1* TG mice using the NucleoSpin RNA Kit (Macherey-Nagel, Düren, Germany). The RNA samples were analyzed with Affymetrix GeneChip mouse genome 430 2.0 Arrays by Takara Bio (Otsu, Shiga, Japan).

### Next-generation sequencing (NGS)

Whole-genome DNA was isolated from *Mlc1* TG mice using the NucleoBond AXG Column (Macherey-Nagel). Identification of the loci of transgene insertion was performed by Takara Bio, followed by NGS on the Illumina sequencing platform.

### qRT-PCR

Total RNA of cells or tissues was extracted using a NucleoSpin RNA kit (Macherey-Nagel). Total RNA extracts were reverse-transcribed using random primers and a QuantiTect Reverse Transcription kit (QIAGEN, Hilden, Germany), according to the manufacturer’s instructions. Real-time PCR was performed using a LightCycler Quick System 350S (Roche Diagnostics), with THUNDERBIRD SYBR qPCR Mix (Toyobo, Osaka, Japan). PCR primers used in this study were as follows: β*-actin* sense primer, 5′-AGCCATGTACGTAGCCATCC-3′; *β-actin* antisense primer, 5’-CTCTCAGCTGTGGTGGTGAA-3’; *Mlc1* sense primer, 5’-CTGACTCAAAGCCCAAGGAC-3’; *Mlc1* antisense primer, 5’-AGCGCAAATAATCCATCTCG-3’; *Mov10l1* sense primer, 5’-TGCTTCTGAACGTGGGACAGG-3’; *Mov10l1* antisense primer, 5’-ACACAGCCAATCAGCACTCTGG-3’; *Ngf* sense primer, 5’-TCAGCATTCCCTTGACACAG-3’; *Ngf* antisense primer, 5’-GTCTGAAGAGGTGGGTGGAG-3’; *Nrf2* sense primer, 5’-GCAACTCCAGAAGGAACAGG-3’; *Nrf2* antisense primer, 5’-GGAATGTCTCTGCCAAAAGC-3’; *Scn9a* sense primer, 5’-AAGGTCCCAAGCCCAGTAGT-3’; *Scn9a* antisense primer, 5’-AGGACTGAAGGGAGACAGCA-3’; *Scn10a* sense primer, 5’-GCCTCAGTTGGACTTGAAGG-3’; *Scn10a*, antisense primer, 5’-AGGGACTGAAGAGCCACAGA-3’; *Trpv1* sense primer, 5’-CCCTCCAGACAGAGACCCTA-3’; *Trpv1* antisense primer, 5’-GACAACAGAGCTGACGGTGA-3’.

### Western blotting

Samples (cells or tissues) were lysed with 10 mM Tris, pH 7.4, containing 150 mM NaCl, 5 mM EDTA, 1% Triton X-100, 1% deoxycholic acid, and 0.1% sodium dodecyl sulfate (SDS). The homogenate was centrifuged at 20,600 *g* for 5 min, and the supernatant was stored at −20°C. Protein concentration was measured using a bicinchoninic acid protein assay kit (Pierce). Equal amounts of protein per lane were electrophoresed on SDS-polyacrylamide gels, and then transferred to a polyvinylidene difluoride membrane. The blots were probed with rabbit anti-SNX25 (1:1000, 13294-1-AP, Proteintech), rabbit anti-TRPV1 (1:100, KM018, Trans Genic), rabbit anti-TrkA (1:10000, ab76291, abcam), goat anti-CD206 (1:1000, AF2535, R&D Systems), rabbit anti-NGF (1:200, sc-548, Santa Cruz Biotechnology), rabbit anti-Nrf2 (1:500, sc-722, Santa Cruz Biotechnology), rabbit anti-HO-1 (1:500, ADI-SPA-896, Enzo Life Sciences, Farmingdale, NY, USA), and rabbit anti-GAPDH (1:2000, ABS16, Burlington, MA, USA) antibodies. Immunoblot analysis was performed with horseradish peroxidase-conjugated anti-mouse and anti-rabbit IgG using enhanced chemiluminescence Western blotting detection reagents (Wako). Data were acquired in arbitrary densitometric units using Scion image software.

### Primary DRG neurons

DRGs from *Snx25* +/− and WT littermate mice were quickly collected in DMEM/F12 medium and incubated for 90 min at 37°C in a 0.2% collagenase solution. After dissociation, DRGs were transferred to a tube containing DMEM/F12 supplemented with 10% fetal bovine serum (FBS), 1% penicillin/streptomycin solution. Ganglia were gently triturated using pipettes. After centrifugation, cells were resuspended in DMEM/F12 supplemented as above and plated on poly-L-lysine-coated culture dishes. Neurons were kept at 37°C in 5% CO_2_ and the medium was changed to DMEM/F12 with B27 supplement 8 h after plating.

### Fluo-4 Calcium imaging

DRG neurons were seeded in 96-well cell culture plates at a density of 1.5 ×☐10^4^ cells per well and cultured overnight. Intracellular calcium responses to capsaicin were measured using Calcium kit II-Fluo4 (CS32, Dojindo, Kumamoto, Japan) in accordance with the manufacturer’s instructions. The temperature of the platform was controlled to 37°C. Cells were fluorescently imaged at 495-nm excitation every 7 s, and the fluorescence intensities of neurons were quantified at 515 nm. Fluorescence intensities of neurons were quantified simultaneously for the entire well. Capsaicin (10 μM) was added to measure the response.

### Bone marrow transplantation (BMT)

BM recipients were male 8-week-old C57BL/6J, *Snx25* +/+, *Snx25* +/−, or *Snx25^loxP/loxP^* mice. Mice were intraperitoneally injected with the chemotherapeutic agent busulfan (30☐μg/g body weight; B2635, Sigma-Aldrich) in a 1:4 solution of dimethyl sulfoxide and PBS 7, 5, and 3☐days prior to bone marrow transfer. All mice were treated with antibiotics (trimethoprim/sulfamethoxazole) for 14☐days after busulfan treatment. Bone marrow-derived cells were obtained from the femur and tibia of 5-week-old C57BL/6-Tg (CAG-EGFP), *Snx25* +/+, *Snx25* +/−, or *Cx3cr1*^*Cre*ERT2/WT^; *Snx25^loxP/loxP^* mice and resuspended in PBS with 2% FBS. Bone marrow-derived cells (1☐×☐10^6^) were transferred to 8-week-old male C57BL/6, *Snx25* +/+, *Snx25* +/−, or *Snx25^loxP/loxP^* recipient mice by tail vein injection (100☐μl). For quantitative analysis, engraftment was verified by determining the percentage of EGFP-expressing cells in the blood. We counted the numbers of EGFP^+^ cells in peripheral blood by flow cytometry and confirmed efficient chimerism as demonstrated by the large proportions of circulating blood leukocytes expressing EGFP.

### Bone marrow-derived macrophage (BMDM) culture

Bone marrow cells were obtained from femur and tibia of 8-week-old male *Snx25* +/+, *Snx25* +/−, or *Cx3cr1*^*Cre*ERT2/WT^; *Snx25^loxP/loxP^* mice and cultured in RPMI-1640 medium containing 10% FBS, 1% penicillin/streptomycin, and 0.01% macrophage colony stimulating factor (M-CSF). After 6 days, the BMDMs were transferred to 3.5-mm dishes in RPMI-1640 containing 10% FBS and 1% penicillin/streptomycin. After overnight incubation, qPCR or Western blot analysis of BMDMs was performed.

### PCR array

The mouse inflammatory response and autoimmunity RT^2^ Profiler PCR Array kit (PAMM-077Z, Qiagen) in a 96-well format was used. This kit profiles the expression of 84 genes that encode inflammatory response, autoimmunity, and other genes related to inflammation. Hind paw skins were quickly dissected 3 d after formalin injection, frozen rapidly, and stored at −80°C until use. Total RNA was purified using the NucleoSpin RNA kit (Macherey-Nagel) in accordance with the manufacturer’s instructions. cDNA was obtained from purified RNA using the RT^2^ First Strand Kit (Qiagen) provided with the PCR Array kit. cDNA template mixed with PCR master mix was dispensed into each well and real-time PCR was performed. Three independent arrays (three animals) were performed.

### Fluorescent in situ hybridization (FISH)

FISH was performed with a probe targeting *Cx3cr1* mRNA using the RNAscope Fluorescent multiplex reagent kit (Advanced Cell Diagnostics, Hayward, CA, USA) according to the manufacturer’s instructions.

### Nerve ligation assay

To assess NGF/TrkA complex trafficking from the periphery toward the DRG cell bodies, we carefully exposed the left sciatic nerve and tightly ligated the nerve with one 6.0 suture in WT and *Snx25* +/− mice. Eight hours after the surgery, mice were terminally anesthetized and quickly perfused with 4% PFA. After perfusion, the left sciatic nerve was excised, post-fixed for 24 h in the same perfusion fixative, cryoprotected in 30% sucrose for 48 h at 4°C, and then frozen in tissue freezing medium (O.C.T.). Longitudinal sections (18 μm) of the left sciatic nerve were cut on a cryostat and then stored at −30°C before staining. Sciatic nerve sections were stained with the primary antibody against rabbit anti-TrkA (1:150, ab76291, abcam). Alexa Fluor 594- (Life Technologies) conjugated IgG was used as the secondary antibody.

### Generation of constructs and transient transfection of 293T cells

PCR cloning was performed to amplify *Snx25* and *Nrf2* cDNA with a primer having an optimal Kozak consensus sequence just before the in-frame first ATG of the mouse *Snx25* and *Nrf2* genes. Fragments were inserted into the pcDNA3.1/Myc-His vector (Invitrogen). Using the LipofectAMINE reagent (Invitrogen), 293T cells were transfected with a *Snx25* and *Nrf2* construct according to the manufacturer’s instructions.

### Quantification and statistical analysis

Quantifications were performed from at least three independent experimental groups. Data are presented as mean ± SEM. Statistical analyses were performed using Student’s *t*-test or Welch’s *t*-test for two groups or one-way ANOVA for multiple groups, and significant differences between group means were identified with the Tukey–Kramer test. Statistical significance is indicated as asterisks. \**p* < 0.05, \*\**p* < 0.01. All n are indicated in figure legends.

